# mRNA cap-binding protein eIF4E1 is a novel regulator of *Toxoplasma gondii* latency

**DOI:** 10.1101/2023.10.09.561274

**Authors:** Michael J. Holmes, Matheus S. Bastos, Vishakha Dey, Vanessa Severo, Ronald C. Wek, William J. Sullivan

## Abstract

The protozoan parasite *Toxoplasma gondii* causes serious opportunistic disease due to its ability to persist in patients as latent tissue cysts. The molecular mechanisms coordinating conversion between proliferative parasites (tachyzoites) and dormant cysts (bradyzoites) are not fully understood. We previously showed that phosphorylation of eIF2α accompanies bradyzoite formation, suggesting that this clinically relevant process involves regulation of mRNA translation. In this study, we investigated the composition and role of eIF4F multi-subunit complexes in translational control. Using CLIPseq, we find that the cap-binding subunit, eIF4E1, localizes to the 5’-end of all tachyzoite mRNAs, many of which show evidence of stemming from heterogenous transcriptional start sites. We further show that eIF4E1 operates as the predominant cap-binding protein in two distinct eIF4F complexes. Using genetic and pharmacological approaches, we found that eIF4E1 deficiency triggers efficient spontaneous formation of bradyzoites without stress induction. Consistent with this result, we also show that stress-induced bradyzoites exhibit reduced eIF4E1 expression. Overall, our findings establish a novel role for eIF4F in translational control required for parasite latency and microbial persistence.

**Significance:** *Toxoplasma gondii* is an opportunistic pathogen of importance to global human and animal health. There are currently no chemotherapies targeting the encysted form of the parasite. Consequently, a better understanding of the mechanisms controlling encystation are required. Here we show that the mRNA cap-binding protein, eIF4E1, is involved in regulating the encystation process. Encysted parasites reduce eIF4E1 levels and depletion of eIF4E1 decreases the translation of ribosome-associated machinery and drives *Toxoplasma* encystation. Together, these data reveal a new layer of mRNA translational control that regulates parasite encystation and latency.

## Introduction

*Toxoplasma gondii* is a widespread protozoan parasite of medical and veterinary importance. *Toxoplasma* can switch from a rapidly growing stage, termed tachyzoite, to a latent tissue cyst stage termed bradyzoite. In newly infected hosts, tachyzoites disperse throughout the body until the immune system mounts a defense which drives bradyzoite formation. Upon immunosuppression, dormant bradyzoites reconvert into proliferative tachyzoites which then cause localized tissue destruction. If untreated, reactivated toxoplasmosis can be fatal.

Given the central role that stage switching plays in *Toxoplasma* pathogenesis and disease, it is important to discern the mechanisms governing this process. As stress has long been known to convert tachyzoites to bradyzoites (1, 2), we initiated studies of stress-induced translational control in *Toxoplasma* (3). The stress-dependent phosphorylation of the alpha subunit of eukaryotic initiation factor-2 (eIF2α) deters the delivery of initiator tRNA to ribosomes, abrogating appropriate start codon recognition and efficiency (4). Consequently, global protein synthesis is curtailed while favoring the selective translation of mRNAs encoding factors that drive an adaptive response. We established that *Toxoplasma* eIF2α phosphorylation accompanies bradyzoite formation (5), as well as transitions in the fellow parasite *Plasmodium falciparum* (6) and the protozoan pathogen *Entamoeba* (7, 8).

Further supporting the role of translational control in parasite differentiation, we conducted a polyribosome profiling study in which we detected the preferential translation of ∼500 mRNAs upon activation of the parasite’s unfolded protein response using a stimulus that promotes bradyzoite formation (5, 9). Notably, translation of *BFD1* mRNA, a MYB-family transcription factor found to be necessary and sufficient for bradyzoite formation (10), was increased 30-fold in our study. Although *BFD1* mRNA is present at equal levels in both tachyzoites and bradyzoites, its protein is only detectable once the parasite begins to differentiate underscoring the crucial role of translational control in parasite life cycle transitions (10-12).

Another major mode of translational control involves the binding of the eIF4F complex to the m^7^G-cap of mRNAs (13). The eIF4F complex, composed of the cap-binding protein eIF4E, the scaffolding protein eIF4G, and the helicase eIF4A, recruits ribosomes and other translation initiation factors, including the multi-subunit eIF3, to the 5’-cap of mRNAs for ribosome scanning and subsequent recognition of start codons. Additionally, the eIF4G component of the eIF4F complex is suggested to engage with the poly(A) binding protein (PABP), creating a closed-loop between the 5’- and 3’-ends of mRNAs that facilitates recycling of translating ribosomes and associated factors (14).

The processes by which eIF4F is modulated for cap-dependent translation can vary among eukaryotic organisms, however this point of control is broadly utilized to regulate gene expression. Examples include mTOR-dependent alterations in eIF4F assembly in animals and the utilization of distinct eIF4F complexes in plants (13, 15). Both mechanisms drive selective mRNA translation by facilitating the preferential loading of translation machinery onto distinct subsets of mRNAs. The role that eIF4F-based translational control plays in *Toxoplasma* gene expression is a critical knowledge gap.

*Toxoplasma* encodes multiple eIF4E and eIF4G paralogs, raising the possibility that heterogeneous eIF4F complexes may permit selective translation in the parasite. Here, we reveal that eIF4E1 facilitates the bulk of translation in tachyzoites. We show that eIF4E1 binds mRNAs at their 5’-ends and associates with two eIF4G paralogs, indicating that distinct eIF4F complexes are used by the parasite to govern protein synthesis. The translation of mRNAs encoding components of the protein synthetic machinery is particularly sensitive to lowered levels of eIF4E1. We discovered that eIF4E1 is a crucial factor required for tachyzoite maintenance; genetic knockdown or chemical inhibition of eIF4E1 causes spontaneous conversion into bradyzoites without the need of stress induction. Together, our findings establish a novel regulatory process critical for differentiation that could serve as a focal point for therapeutics.

## Results

### Functional profiling of eIF4E paralogs in tachyzoites

Based on IF4E domain homology searches (PFAM: PF01652 from toxodb.org) *Toxoplasma* encodes three members of the eIF4E family, designated eIF4E1, -2, and -3 (Fig. 1A) (16). eIF4E1 (TGME49_223410) and eIF4E2 (TGME49_315150) have key features of the eIF4E domain that are shared among other studied model organisms (Fig. S1A-B). eIF4E1 possesses all residues predicted to be required for binding the m^7^G cap, encodes the S/TVxxF eIF4G binding motif, and is 26 kDa, a typical size for eIF4E family proteins. In contrast, eIF4E2 is 50 kDa due to long N- and C-terminal extensions and an insert between its second and third conserved aromatic residues. In addition, several conserved aromatic resides lack conservation including near its eIF4G binding motif. The eIF4E3 homolog (TGME49_312560) also has noncanonical features. In addition to being very large (214 kDa), its IF4E domain displays degeneracy in some conserved aromatic residues and within its eIF4G-binding motif (Fig. S1A).

**Figure 1.**
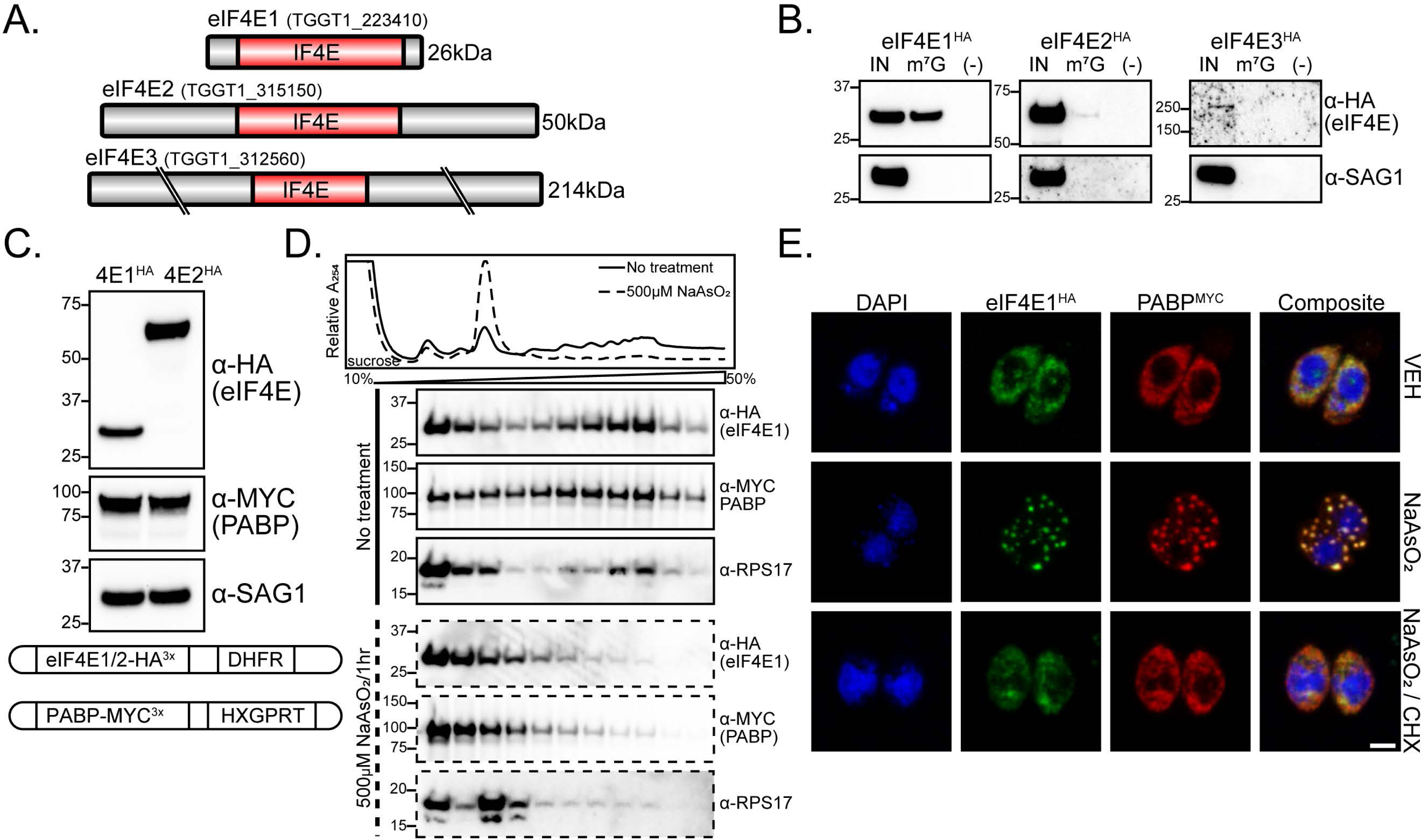
Functional profiling of eIF4E paralogs. A) IF4E domain organization of *Toxoplasma* eIF4E paralogs. Gene accession numbers and expected molecular weight are indicated. All cartoons are drawn to scale. B) Affinity purification of eIF4E^HA^ paralogs with m^7^G-functionalized resin. Tachyzoite lysate was incubated with m^7^G-functionalized or control resin and subjected to western blot analysis. SAG1 was included as control for non-specific binding of lysate to the resin. Molecular weight markers for each blot are noted in kDa. C) Generation of dual-tagged lines by endogenously tagging PABP with MYC epitope in HA-tagged eIF4E lines. Western blot showing relative abundance of eIF4E1^HA^ and eIF4E2^HA^ along with PABP^MYC^ and SAG1 loading control. A cartoon summarizing genetic manipulations is displayed below the blot. D) Analysis of eIF4E1^HA^ and PABP^MYC^ sedimentation by polysome profiling of tachyzoites exposed to oxidative stress induced by incubation with 500 µM NaAsO_2_ for 1 h or no treatment. The trace is generated by continuous absorbance at A254 which measures RNA content with major peaks at rRNA subunits, monosome, and heavy polysomes. Fractions of equal volumes were collected from the gradient and proteins were enriched by TCA precipitation and analyzed by western blot. An antibody directed against RPS17 that is suggested to recognize both *Toxoplasma* and human orthologs was included on the blot as a reference marker for protein sedimentation profiles. E) Analysis of eIF4E1 localization to stress granules upon oxidative stress. eIF4E1^HA^ and PABP1^MYC^ were visualized by immunofluorescence microscopy and DNA by DAPI staining. Samples were treated with 500 µM NaAsO_2_ ± 100 µg/ml cycloheximide (CHX) for 1 h prior to fixation. The translation elongation inhibitor CHX was included as a control to prevent stress granule formation. Microscopy scale bar = 2 µm.

To address the functional properties of the eIF4E proteins in *Toxoplasma*, we engineered three RH strain parasite lines in which each respective eIF4E was endogenously tagged with HA epitope at its C-terminus. The tagged proteins displayed the predicted size as indicated by western blot analysis (Fig. S1C). As expected for putative translational machinery, each eIF4E family member was localized exclusively to the parasite cytoplasm, although eIF4E3 expression was very low (Fig. S1D). We next tested whether each family member could bind the m^7^G mRNA cap by incubating tachyzoite lysate with m^7^G-functionalized agarose resin. Only eIF4E1^HA^ bound the functionalized resin with high affinity (Fig. 1B). Due to its degeneracy, atypically large size, and very low expression profile, eIF4E3 was excluded from further analysis and we focused our studies on eIF4E1 and eIF4E2.

To compare eIF4E1 and eIF4E2 functions, we generated dual-tagged lines by fusing a C-terminal MYC tag to PABP (TGME49_224850; Fig. 1C). Since both paralogs were labeled with the same epitope tag, we were able to determine that eIF4E2^HA^ is more abundant than eIF4E1^HA^ (Fig. 1C). Given that eIF4E2^HA^ bound m^7^G resin with little to no affinity despite being more abundant, we conclude that the eIF4E1^HA^ ortholog is the primary cap-binding protein in tachyzoites.

To address whether eIF4E1 and eIF4E2 share other features of documented cap-binding proteins, we first measured eIF4E1^HA^ association with translating mRNAs by polysome profiling. Under non-stressed conditions in tachyzoites, eIF4E1^HA^ and PABP^MYC^ were present within the heavy polysomes in the gradient, which represents mRNAs associated with multiple translating ribosomes (Fig. 1D). Both tagged proteins shift to non-translating fractions when challenged with sodium arsenite, a potent trigger of oxidative stress that sharply lowers translation initiation and shifts heavy polysomes to monosomes and free ribosomal subunits (17) (Fig. 1D). In other eukaryotes, the acute inhibition of translation initiation by arsenite also causes aggregation of translational initiation machinery along with general factors such as PABP into stress granules (18). Consistent with this idea, we found that eIF4E1^HA^ localizes to stress granules along with PABP^MYC^ in response to arsenite (Fig. 1E). We performed similar experiments with eIF4E2^HA^ and found that it also dissociates from heavy polysomes and redistributes to stress granules upon treatment with arsenite (Fig. S2A-B). These results indicate that although only eIF4E1 is associated with the m^7^G 5’-cap structure, both eIF4E1 and eIF4E2 interact with translating mRNAs.

### eIF4E1 interacts with two eIF4G paralogs

We next addressed the protein-binding partners for eIF4E1 and eIF4E2. HA-tagged eIF4E paralogs were immunoprecipitated from tachyzoites under control and sodium arsenite-treated conditions. Untagged parental parasites were used as a negative control to screen for non-specific interactions. We found that eIF4E1^HA^ interacted with eIF4G1 and eIF4G2, along with all but one subunit of the eIF3 complex, PABP, and eIF4A (Fig. 2A and Table S1). Given the multi-domain organization of the eIF4G family and their role in recruiting eIF4A, PABP, and eIF3, we hypothesize that many of these proteins co-purified with eIF4E1^HA^ through secondary contact sites. By contrast, eIF4E2^HA^ did not associate with any eIF4F-related components or other initiation factors; its top interacting protein, the hypothetical protein TGGT1_244460, is of unknown function but encodes four copies of the putative eIF4E-interacting motif YXXXXLΦ (Fig. S2C, Table S1).

**Figure 2.**
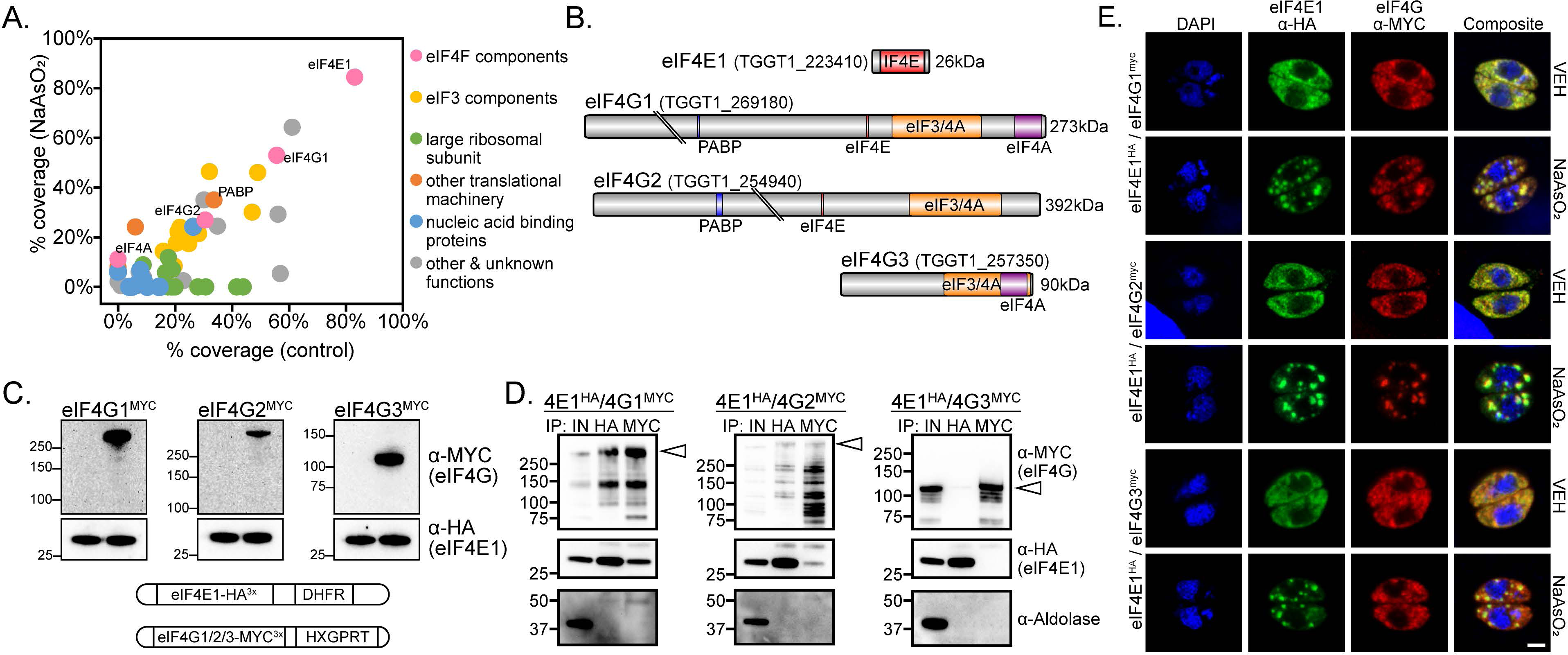
eIF4E1 binds two eIF4G paralogs. A) eIF4E1^HA^ interactomes under stressed (y-axis; 500 µM NaAsO_2_ for 1 h) or non-stressed (x-axis) tachyzoites. Affinity purification of eIF4E1^HA^ was conducted by immunoprecipitation with α-HA magnetic beads followed by mass spectrometry identification from two biological replicates. Interacting proteins that were absent in the untagged control immunoprecipitations and not localized to *Toxoplasma* secretory organelles (42) are displayed and colored by functional identity. The average percentage of coverage from each identified protein is displayed. B) Domain and motif architecture of the three eIF4G paralogs encoded by *Toxoplasma*. Gene accession numbers and expected molecular weight are indicated. The location of protein-interacting domains and motifs were determined using alignments of the *Toxoplasma* proteins compared to the domains of Human eIF4G1 available on UniProt (Q04637). All cartoons are drawn to scale. C) Generation of dual-tagged lines by endogenously tagging eIF4G paralogs with MYC epitope in eIF4E1^HA^ background. The first lane of each western blot is the parental (eIF4E1^HA^) line. Each eIF4G^MYC^ paralog displayed the predicted molecular weight. The blots were probed with αHA as a loading control. Molecular weight markers for each blot are noted in kDa. A cartoon summarizing genetic manipulations is displayed below the blot. D) Reciprocal co-immunoprecipitation of eIF4E1^HA^ and eIF4G1/2/3^MYC^. The first lane of each blot presents total protein lysate. The blot was probed with α-aldolase as a control for non-specific protein binding to the beads. A white triangle appears on each blot to indicate the expected molecular weight of each intact eIF4G paralog. The milder lysis conditions required for co-immunoprecipitation, resulting in multiple lower molecular weight products for each eIF4G paralog, is characteristic of post-lysis protein degradation. E) Analysis of eIF4G^MYC^ paralog localization to stress granules during oxidative stress. eIF4E1^HA^ and eIF4G^MYC^ paralogs were visualized by immunofluorescence microscopy and DNA by DAPI staining. Samples were treated with 500 µM NaAsO_2_ ± 100 µg/ml cycloheximide (CHX) for 1 h prior to fixation. The translation elongation inhibitor CHX was included as a control to prevent stress granule formation. Microscopy scale bar = 2 µm.

In *Toxoplasma*, eIF4G1 and eIF4G2 are both long paralogs that include the conserved YXXXXLΦ amino acid motif that interacts with eIF4E (Fig. 2B) (19). In contrast, eIF4G3 resembles an N-terminally truncated eIF4G paralog, which is present in some organisms and is missing the eIF4E-interacting motif (Fig. 2B) (20). To further validate the putative interactions between eIF4E1 and eIF4G paralogs, we MYC-tagged each eIF4G paralog at its C-terminus in parasites expressing eIF4E1^HA^ (Fig. 2C). Each protein in these dual-tagged lines displayed the predicted molecular weights as indicated by western blot analyses (Fig. 2C). We then conducted reciprocal co-immunoprecipitations using these parasite lines. Consistent with our unbiased proteomic method, affinity purification of eIF4E1^HA^ recovered eIF4G1^MYC^ and eIF4G2^MYC^ but not eIF4G3^MYC^ (Fig. 2D). Pulldown of eIF4G1^MYC^ and eIF4G2^MYC^ also showed association with eIF4E1^HA^. The milder lysis conditions that were required to maintain the eIF4E-eIF4G interactions resulted in partial degradation of the eIF4G orthologs despite the addition of protease and phosphatase inhibitor cocktail to the reaction; arrowheads denote the full-length products (Fig. 2D). Finally, we assessed whether the eIF4Gs were present in stress granules upon sodium arsenite-induced suppression of translation. Consistent with their being interacting partners, we found that eIF4G1^MYC^ and eIF4G2^MYC^ localized to stress granules whereas eIF4G3^MYC^ did not, suggesting the latter may have a different function (Fig. 2E). These results suggest that *Toxoplasma* eIF4E1 can form heterogenous complexes with eIF4G1 or eIF4G2.

### CLIPseq reveals that eIF4E1 binds the 5’-end of all tachyzoite mRNAs

The 5’-end of eukaryotic mRNAs have an m^7^G cap that is recognized by eIF4E and related proteins. Using our HA-tagged eIF4E lines, we conducted CLIPseq (21) in tachyzoites to identify which mRNA subsets are bound by each paralog. The mRNA regions that were enriched for eIF4E1^HA^ or eIF4E2^HA^ binding are reported in Table S2. Consistent with our m^7^G-affinity purification data (Fig. 1B), eIF4E2^HA^ was not enriched at the 5’-ends of mRNAs (Fig. S2D). Instead, eIF4E2^HA^ was more generally enriched throughout the non-coding mRNA regions. By contrast, eIF4E1^HA^ was enriched at the transcriptional start site (TSS) of every transcribed protein coding mRNA expressed in tachyzoites (Fig. 3A-B). Given the pervasive association of eIF4E1^HA^ with TSSs, which represent the 5’-ends of the encoded mRNAs, we conclude that it plays the predominant role in coordinating cap-dependent translation in this life cycle stage.

**Figure 3.**
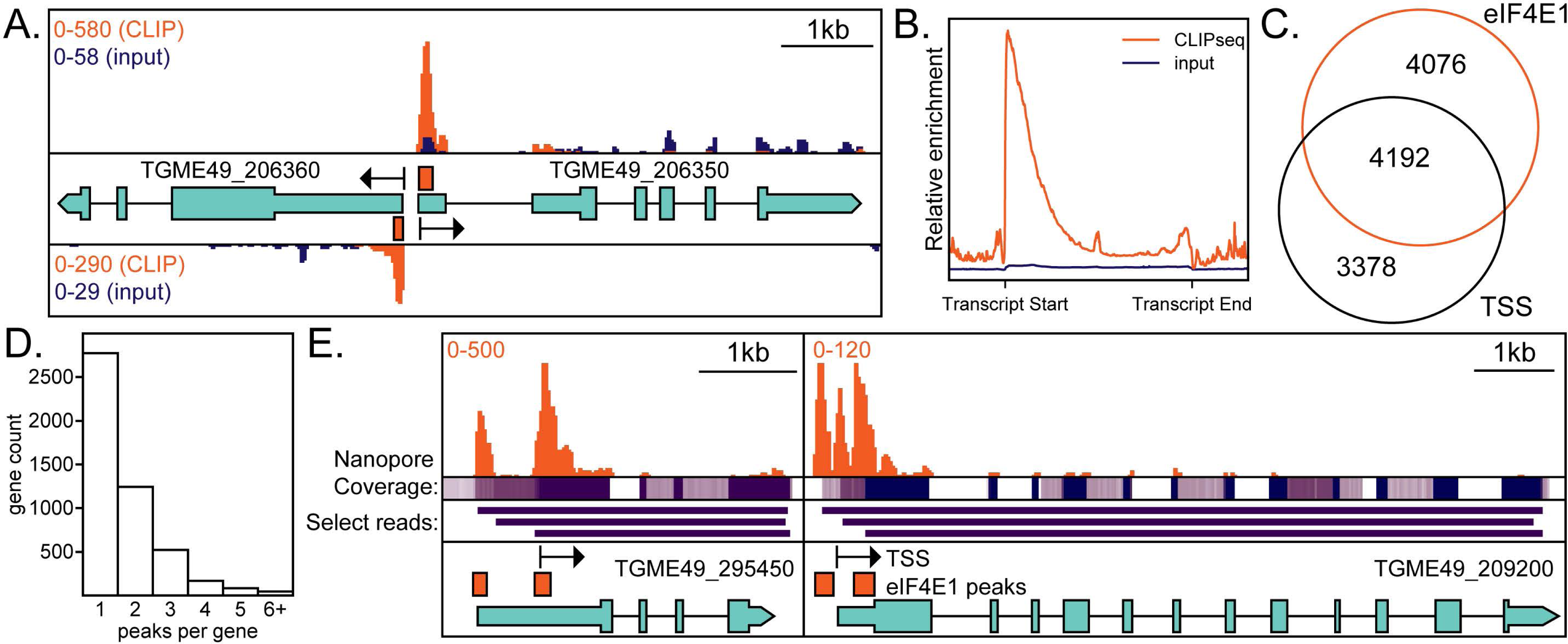
eIF4E1^HA^ associates with the 5’ end of mRNAs. A) An example of a bidirectional promoter demonstrates the specificity of interaction between eIF4E1^HA^ and m^7^G-capped mRNAs. Sequencing tracks from size-matched input samples (input) are shown for reference. The scale in the top left side of the panel indicates the relative abundance of CLIPseq (orange) and input reads (blue). Black arrows indicate annotated transcriptional start sites (TSS) as determined by RAMPAGEseq (22). Orange bars are regions statistically enriched by eIF4E1^HA^ CLIPseq. Relevant gene accession numbers are indicated. B) A metagene plot demonstrating CLIPseq enrichment over input and that eIF4E1^HA^ is associated with the TSS. C) Analysis of the proximity (within 20nt) of eIF4E1^HA^-enriched sites to annotated TSSs. D). Analysis of the number of eIF4E1^HA^-enriched regions per gene. E) Two examples of alternate TSS usage. eIF4E1^HA^-enriched regions are supported by nanopore reads as evidenced by coverage and select individual read tracks, displayed in purple (24). The scale in the top left side of the panel indicates the relative abundance of CLIPseq reads (orange). Relevant gene accession numbers are indicated.

To further address the role of eIF4E1 in mRNA translation, we used our CLIPseq data to quantitate the degree of eIF4E1^HA^ engagement with mRNAs and compared that to total mRNA abundance (Fig. S3A). This latter dataset was obtained from a ribosome profiling experiment that we conducted as a part of this study (described below). We determined a strong correlation between mRNA abundance and its engagement with eIF4E1^HA^ (Fig. S3A). The relationship between eIF4E1^HA^ association and mRNA abundance was not observed for genes encoded in the apicoplast, which is consistent with the idea that this organelle follows a bacterial-like translational paradigm that does not employ cap-binding proteins. Our results support a general function for eIF4E1 in the initiation of *Toxoplasma* translation since its engagement with mRNAs is correlated with mRNA abundance with little evidence for transcript specificity.

We compared our eIF4E1^HA^ CLIPseq results to the annotated TSSs that were generated by RAMPAGEseq and appear on toxodb.org (22). Most eIF4E1^HA^-enriched regions and annotated TSSs fell within 20nt of each other, which highlights the robustness of the experiment (Fig. 3C). We quantified the number of eIF4E1^HA^-associated peaks per gene (Fig. 3D). While most genes had a single statistically enriched peak, many were associated with multiple eIF4E1^HA^ binding sites, which is indicative of TSS heterogeneity for these genes. Illustrations of multiple TSS usage are shown in Fig. 3E. For example, the SSNA1/DIP13 gene (23), encoded by TGME49_295450, displayed two eIF4E1^HA^ binding sites, one of which has been previously suggested by RAMPAGEseq (22). The two eIF4E1^HA^ peaks are a likely indication of alternative transcription initiation sites, and both are further supported by nanopore sequencing reads available through toxodb.org (16, 24). In another example, the hypothetical protein encoded by TGME49_209200 has three eIF4E1^HA^-binding sites, each also supported by nanopore sequencing (24). These results show that eIF4E1 binds globally to the 5’-ends of mRNAs and this association can be applied for curation of multiple gene TSSs.

### Depletion of eIF4E1 reduces translation and replication

To address the contribution of eIF4E1 in parasite viability and translation in tachyzoites, we endogenously tagged eIF4E1 with an HA epitope and a minimal auxin inducible degron (mAID) for conditional depletion in RH strain parasites (25). eIF4E1^mAID-HA^ was efficiently degraded, with minimal detection of the protein as indicated by western blot following 2 h treatment with 500 μM of the auxin 3-indoleacetic acid (IAA; Fig. 4A). Using this targeted depletion strategy, we conducted a plaque assay to determine the contribution of eIF4E1 to parasite growth over the course of multiple lytic cycles. A complete absence of plaques was observed when eIF4E1^mAID-HA^ was depleted for a week (Fig. 4B). Interestingly, a severe, but not complete, loss of plaquing was seen when IAA was only added as a 24 h pulse (Fig. 4B), indicating that the parasites were able to recover from transient eIF4E1^mAID-HA^ depletion. Parasite replication was also reduced, but not eliminated, over a 16 h IAA treatment (Fig. 4C). Similar experiments conducted on eIF4E2^mAID-HA^ tachyzoites revealed that its depletion has no effect on parasite replication, indicating that it is not essential for tachyzoites viability (Fig. S2E-G).

**Figure 4.**
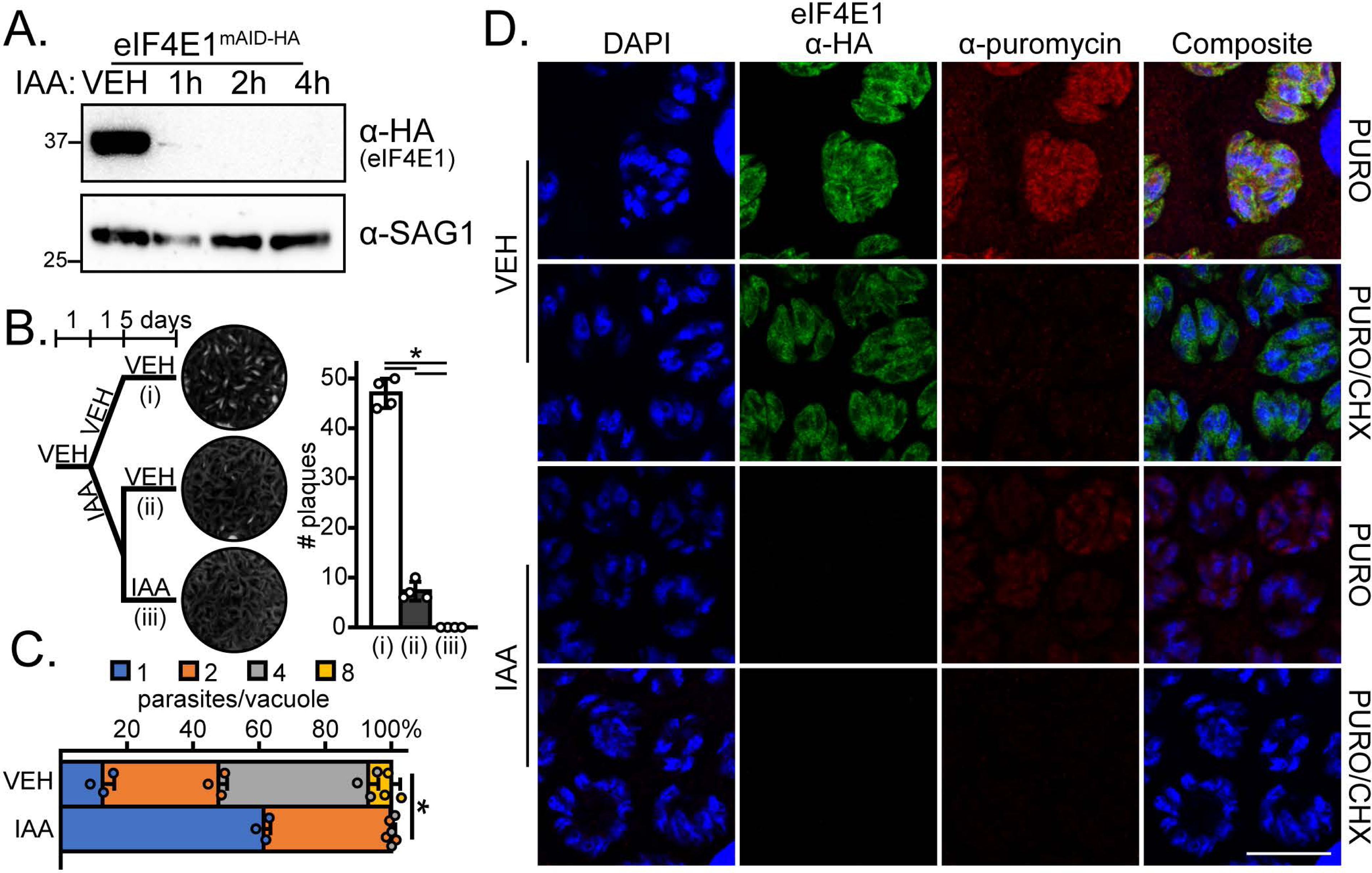
eIF4E1^mAID-HA^ integrity is required for efficient tachyzoite replication and translation. A) Analysis of eIF4E1^mAID-HA^ depletion over time upon addition of 500 µM IAA by western blot. The blot was probed with α-SAG1 as a loading control. Molecular weights are indicated in kDa. B) Plaque assay of vehicle (0.5% DMSO)-treated parasites, parasites pulsed for 24 h with 500 µM IAA or parasites treated with IAA for 6 days. Mean plaque numbers ± standard deviation were tested for statistical significance by a one-way ANOVA followed by a Student’s t-test assuming unequal variances. *p ≤ 0.01. C) Analysis of parasite replication after a 16 h treatment with 500 µM IAA. Mean number of parasites per vacuole with standard deviation is shown. Statistical significance of the change between the mean number of parasites per vacuole was determined by Student’s t-test assuming unequal variances. *p ≤ 0.01. D) Representative immunofluorescence assay images of eIF4E1^mAID-HA^ parasites treated with either DMSO vehicle or with IAA for 4 h. A 10 min pulse with 10 µg/ml puromycin to label newly synthesis proteins was included. eIF4E1^mAID-HA^ and puromycinylated peptides were visualized by immunofluorescence microscopy and DNA by DAPI staining. The translation elongation inhibitor CHX was added concurrently with puromycin to completely block protein synthesis as a control. Microscopy scale bar = 10 µm.

Multiple lines of research involving model organisms have suggested that the translation of some mRNAs is more sensitive to eIF4E1 levels than others (4, 13, 19). To address whether eIF4E1 imparts any selectivity in *Toxoplasma* mRNA translation, we first verified that eIF4E1^mAID-HA^ depletion reduced global translation as judged by the incorporation of puromycin into nascently translated proteins (Fig. 4D). Next, we conducted a ribosome profiling (RIBOseq) experiment in tachyzoites treated with IAA for 4 h to assess the consequences of eIF4E1^mAID-HA^ depletion (Table S3). The amount of ribosome protected footprints (RPF) in DMSO-treated (control) parasites strongly correlated with mRNA abundance, again supporting the general function of eIF4E1 in directing cap-dependent translation in tachyzoites (Fig. S3B).

We observed a generalized decrease in transcript abundance and mRNA translation upon eIF4E1^mAID-HA^ depletion (Fig. 5A-B). While only 41 and 54 genes were upregulated at the mRNA and RFP levels, respectively, 223 genes were downregulated at the mRNA level and 525 genes demonstrated decreased translation (Table S3). Gene ontology analysis revealed that translationally downregulated genes were strongly enriched for ribosome-associated machinery (Fig. 5C-D). We found a decrease in the translational efficiency of 153 genes which were also enriched for ribosomal-associated machinery (Fig. 5E-F). We examined the characteristics of *Toxoplasma* 5’-leaders and found that those genes whose translational efficiency were sensitive to eIF4E1 depletion tended to have much shorter 5’-leaders than the average transcript (Fig. 5G). The translationally repressed genes were not clustered according to their transcript abundance or eIF4E engagement (Fig. S3) and they shared the same pyrimidine-rich motif at their transcriptional start site as the total transcriptome (Fig. 5H).

**Figure 5.**
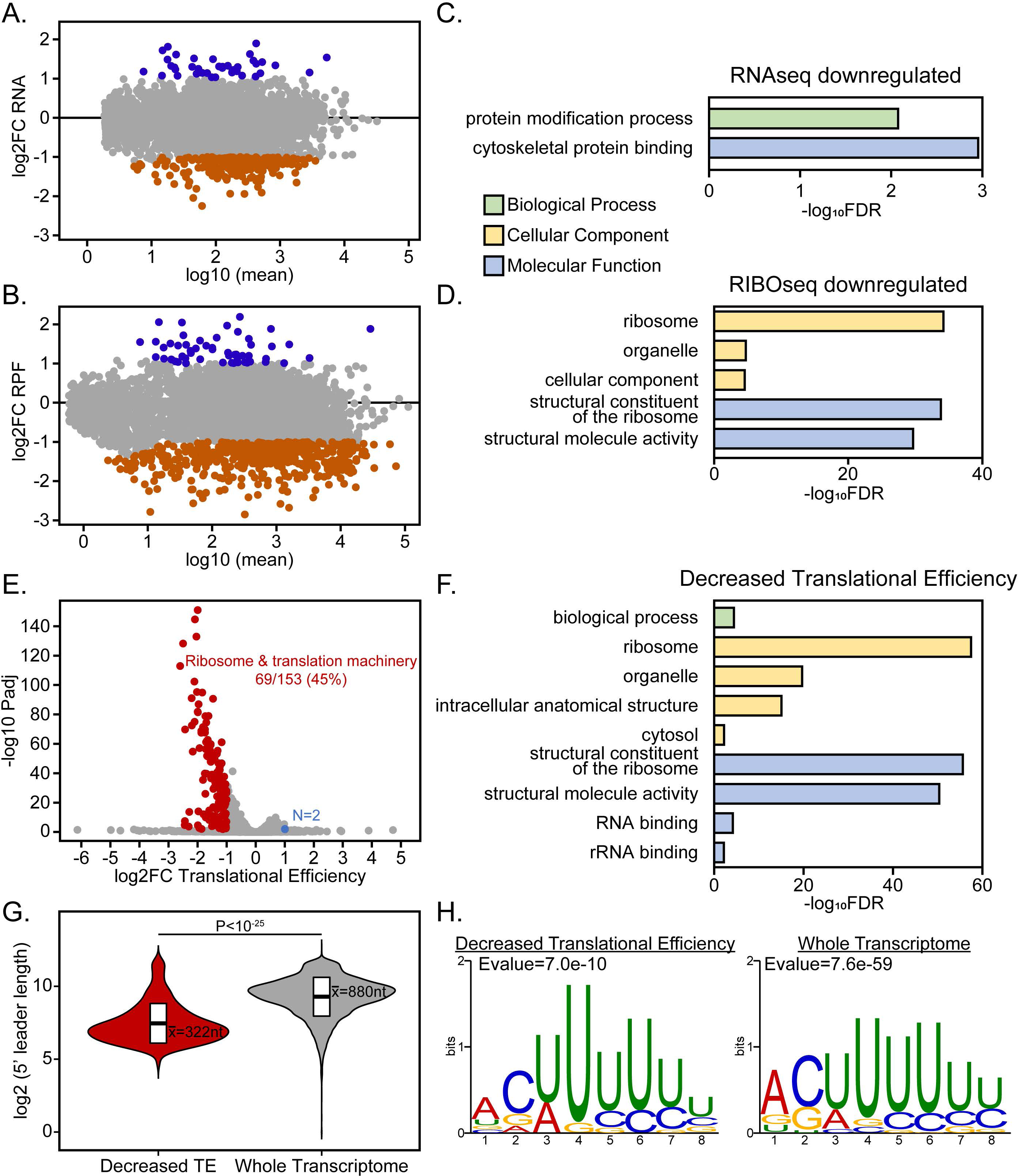
eIF4E1-depleted tachyzoites display reduced translational efficiency of ribosome-associated proteins. A-B) MA plots showing changes in (A) mRNA abundance and (B) ribosome protected footprints (RPF) upon depletion of eIF4E1^mAID-HA^ for 4 h with 500 µM IAA. C-D) Gene ontology enrichment categories of (C) mRNA and (D) RPF changes caused by eIF4E1^mAID-HA^ depletion. The x-axis displays the log_10_ transformed Bonferroni-adjusted false discovery rate with a cutoff of Padj ≤ 0.01. E) eIF4E1^mAID-HA^-dependent changes in translational efficiency. F) Gene ontology enrichment categories of genes that are less efficiently translated upon eIF4E1^mAID-HA^ depletion. G) Analysis of 5’-leader length for those genes whose translational efficiency is dependent on eIF4E1 levels compared to the general *Toxoplasma* transcriptome. H) Motif analysis near the transcriptional start sites of genes categorized by translational efficiency.

Although there was no significant Gene Ontology enrichment seen for the upregulated gene set upon eIF4E1^mAID-HA^ depletion, we did find that two AP2 family transcription factors, AP2IX-9 and AP2IV-3, were upregulated at the mRNA and RPF levels (Table S3). These two transcription factors have been implicated in promoting bradyzoite formation (26, 27), suggesting a linkage between eIF4E1-dependent translation and bradyzoite formation in *Toxoplasma*.

### Depletion of eIF4E1 drives the formation of bradyzoites

In addition to the slow growth phenotype that we observed upon eIF4E1 depletion (Fig. 4B-C), our RIBOseq analysis suggested that loss of eIF4E1 can trigger expression of genes contributing to bradyzoite differentiation (Table S3). Therefore, we addressed whether eIF4E1 is involved in this stage transition. Indeed, depletion of eIF4E1^mAID-HA^ led to the formation of bradyzoite-containing cysts in the absence of stressful culture conditions as evidenced by *Dolichos biflorus* lectin (DBL) staining (Fig. 6A). We quantified the rate of bradyzoite conversion and found that depletion of eIF4E1^mAID-HA^ was more effective than the current gold-standard of stress-induced conversion (combined alkaline pH and CO_2_ depletion; Fig. 6A-B). Consistent for a type I RH parental strain, stress-induced bradyzoite formation was inefficient based on the high degree of host lysis after 5 days (Fig. 6B). In striking contrast, no host cell lysis was observed upon IAA-induced eIF4E1^mAID-HA^ depletion which is unusual for RH strain parasites that are known to be largely refractory to cyst formation (28). Importantly, those ∼50% of vacuoles that were not fully DBL-positive displayed partial staining along their periphery suggesting they were in the process of transitioning to bradyzoites. We excluded IAA itself as a trigger for bradyzoite formation since IAA-induced depletion of eIF4E2^mAID-HA^ did not impact plaquing efficiency compared to a vehicle control (Fig. S2F).

**Figure 6.**
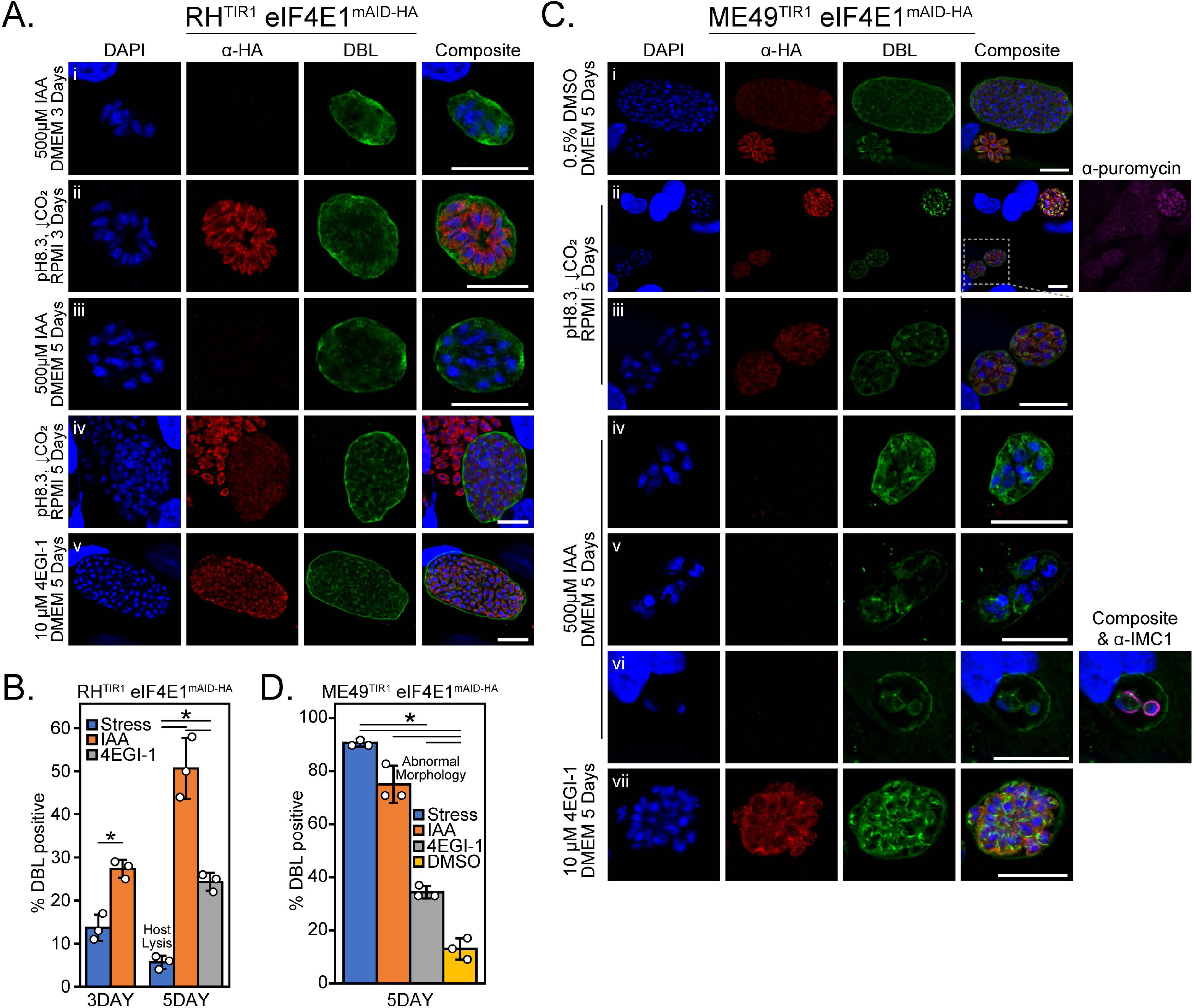
Prolonged depletion of eIF4E1 induces bradyzoite formation. A) Representative immunofluorescent microscopy images of RH strain eIF4E1^mAID-HA^ parasites after 3- or 5-days treatment with alkaline media to for stress-induced bradyzoite formation (panels i, iii), after incubation with 500 µM IAA (panels ii, iv), or incubation with 10 µM 4EGI-1 (panel v). eIF4E1^mAID-HA^ was visualized by immunofluorescence microscopy and staining with FITC-conjugated *Dolichos biflorus* lectin (DBL) was included as a marker of the cyst wall along with DAPI staining for DNA. Microscopy scale bar = 10 µm. B) Quantitation of vacuoles that were positive for a contiguous cyst wall around the entirety of the vacuole. The widespread host cell lysis seen after 5 days treatment under alkaline stress is noted denoted on the chart. The rates of cyst formation (standard deviation is shown) were tested for statistical significance by a one-way ANOVA followed by a Student’s t-test assuming unequal variances. *p ≤ 0.01. C) Representative immunofluorescence microscopy images of ME49 strain eIF4E1^mAID-HA^ parasites after treatment for 5 days in the conditions outlined in (A). A 0.5% DMSO vehicle treated condition (panel i) was included to determine the rate of spontaneous differentiation. A representative image of stress-induced bradyzoites (panel ii) is included. A comparison of the relative translational capacity of tachyzoites (DBL negative vacuole, upper right) and bradyzoites (DBL positive vacuoles, lower left) as determined by puromycin labeling of nascent peptides is shown on the far right of panel ii. An enlarged inset of the DBL-positive vacuoles (panel iii) is displayed to better see the morphology of the vacuole contents. Representative images of IAA-treated DBL-positive vacuoles with normal (panel iv) and abnormal morphology (panels v-vi), as shown incomplete DBL staining around its circumference, parasites are shown. The morphology of individual parasites is revealed by IMC1 staining on the far right of panel vi. Representative image of parasites incubated with 10 µM 4EGI-1 for five days is also included (panel vii). Microscopy scale bar = 10 µm. D) Quantitation of vacuoles that were positive for a contiguous cyst wall around the entirety of the vacuole. The proportion of vacuoles with abnormal morphology were not counted as DBL-positive. The rates of cyst formation (standard deviation is shown) were tested for statistical significance by a one-way ANOVA followed by Student’s t-test assuming unequal variances. *p ≤ 0.01.

Stress-induced bradyzoite cysts displayed reduced eIF4E1 staining compared to adjacent undifferentiated tachyzoites (Fig. 6A, panel iv), demonstrating that lowered eIF4E1 abundance accompanies bradyzoite formation. To determine whether a reduction in the assembly of eIF4F complexes also promotes bradyzoite formation, we utilized a small molecule inhibitor of the eIF4E-eIF4G interaction called 4EGI-1 (29). We determined that 4EGI-1 had an anti-replicative effect on *Toxoplasma* at 10 µM but had minimal effect on the confluent human fibroblast host cells at this concentration (Fig. S4A). Parasites treated with 10 µM 4EGI-1 formed bradyzoites (Fig. 6A-B), suggesting that interfering with the assembly of eIF4F complexes triggers *Toxoplasma* differentiation.

We next engineered the expression of eIF4E1^mAID-HA^ in the type II ME49 background to determine whether our observations from RH parasites can be extended to a cystogenic strain that readily forms bradyzoites (Fig. S5A-B). As for type I parasites, DBL-positive cysts expressed eIF4E1^mAID-HA^ at a reduced level compared to adjacent undifferentiated vacuoles (Fig. 6C, panels i-ii). Furthermore, puromycin labeling revealed that global translation was reduced in bradyzoites compared to tachyzoites (Fig. 6C, panel ii), consistent with lowered eIF4E1 levels during differentiation.

As seen for RH parasites, eIF4E1^mAID-HA^ depletion or treatment with 4EGI-1 also induced bradyzoite formation in the ME49 background (Fig. 6C, panel iv and vii). Treating ME49 parasites with IAA, 4EGI-1, or alkaline media triggered bradyzoite formation at a greater rate than that observed in vehicle (DMSO) controls. These findings indicate that eIF4E1mAID-HA depletion, impairment of eIF4F assembly, or stress all promoted bradyzoite formation as it did for the RH strain (Fig. 6D). Interestingly, eIF4E1^mAID-HA^ depletion in the ME49 strain caused morphological defects in ∼25% of vacuoles (Fig. 6C-D). These partially DBL-positive vacuoles were loosely packed with misshaped parasites as viewed by IMC1 staining (Fig. 6C, panels v and vi), suggesting that loss of eIF4E1 may eventually be lethal.

To evaluate whether the bradyzoites formed due to the loss of eIF4E^1mAID-HA^ were viable, we monitored their ability to reconvert to tachyzoites. RH strain parasites were able to recover from 6 days after loss of eIF4E1^mAID-HA^ whereas ME49 were not, even when these type II parasites were left to recover for two weeks longer than their paired stress-induced treatment (Fig. 7A-B, and data not shown). RH and ME49 tachyzoites express eIF4E1 at similar levels, suggesting that the difference in reactivation into tachyzoites is not due to a strain-specific abundance of initial eIF4E1 protein or residual protein after IAA addition (Fig. 7C). Considered together, these results are the first to demonstrate a novel role for the eIF4F complexes in maintaining the tachyzoite stage as depletion of eIF4E1 provokes robust differentiation into bradyzoites.

**Figure 7.**
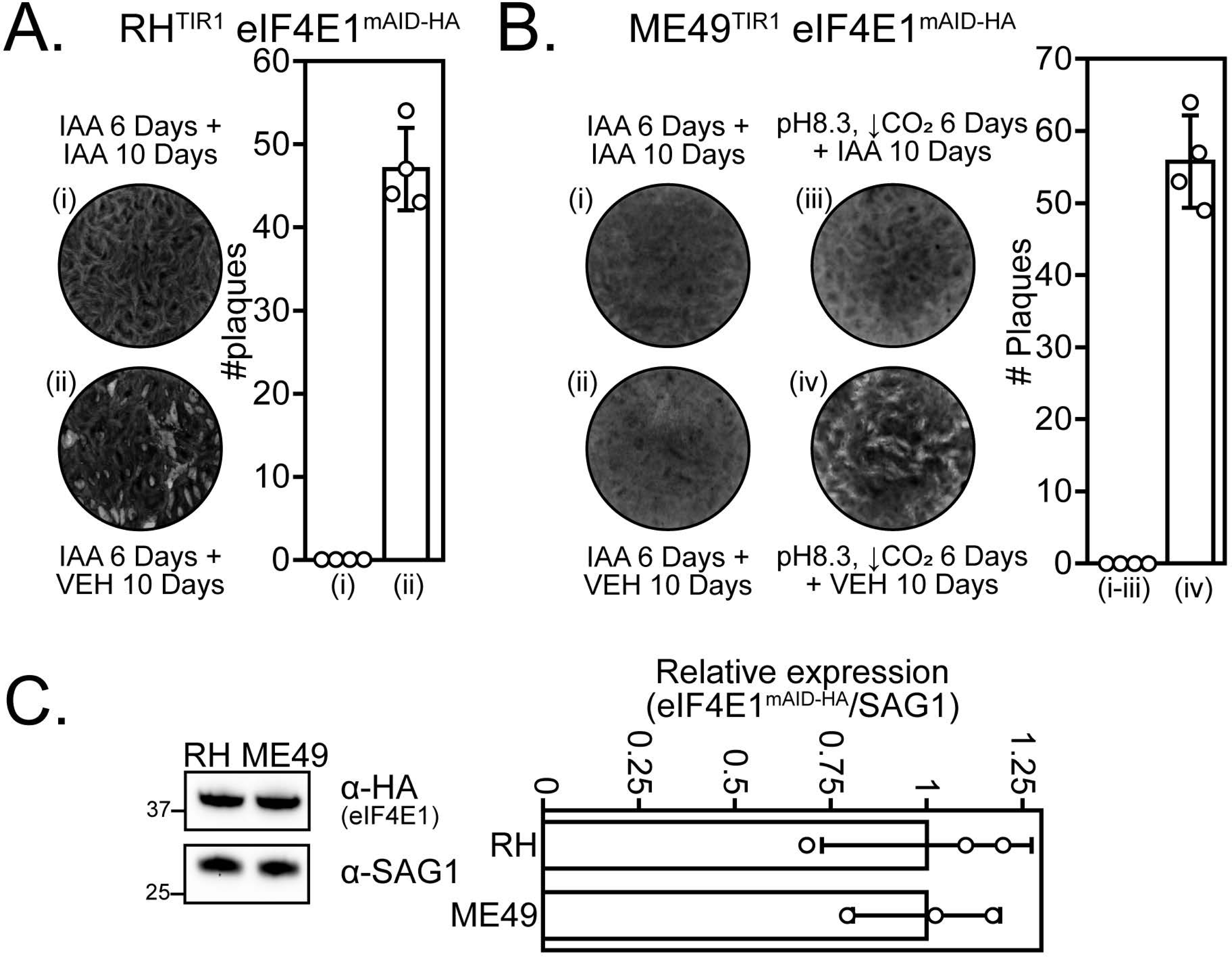
Reactivation assay of eIF4E1^mAID-HA^ parasites after stress-induced or IAA-induced bradyzoite formation. A) RH eIF4E1^mAID-HA^ strain parasites were treated for 6 days with 500 µM IAA to induce bradyzoite formation; cultures were then allowed to recover in 0.5% DMSO (vehicle) or left in IAA for 10 days before assessing parasite growth by plaque assay. A bar graph shows quantitation of the plaque number ± standard deviation from four replicates. B) ME49 eIF4E1^mAID-HA^ strain parasites were treated for 6 days with 500 µM IAA to induce bradyzoite formation then allowed to recover in 0.5% DMSO vehicle or left in IAA for 10 days before assessing parasite growth by plaque assay. An alkaline stress treatment without IAA was included to assess the general ability of parasites to recover from stress-induced differentiation. A bar graph shows quantitation of the plaque number ± standard deviation from four replicates. C) Western blot analysis to determine the relative expression levels of eIF4E1^mAID-HA^ in RH and ME49 strains. Molecular weight markers are shown in kDa. Densitometry of the relative eIF4E1^mAID-HA^ expression ± standard deviation compared to the SAG1 loading control was conducted and quantitated from three biological replicates.

## Discussion

We found that short-term depletion of eIF4E1 reduced translation and delayed replication (Fig. 4). Closer analysis revealed that parasites lacking eIF4E1 were spontaneously converting to bradyzoites at high frequencies over a matter of days, which was unexpected given that our initial studies were conducted in RH strain parasites that are widely considered to be non-cystogenic (Fig. 6). Depletion of eIF4E1 thus offers a new means to induce bradyzoite formation in vitro in the absence of exogenous stress. The ability of small molecule 4EGI-1, which impairs the binding of eIF4G to eIF4E (29), along with the observed decrease of eIF4E1 expression in stress-induced or spontaneously generated bradyzoites, points to a conserved strategy of modulating eIF4F levels to coordinate bradyzoite formation.

### Profiling the eIF4E mRNA cap-binding family in *Toxoplasma*

*Toxoplasma* encodes three eIF4E-like proteins, with only one – eIF4E1 – binding to m^7^G functionalized resin and localizing to the 5’-ends of tachyzoite mRNAs. Here, we applied CLIPseq to determine that eIF4E1 engages with the 5’-end of every transcribed mRNA in tachyzoites and to profile mRNA 5’-end heterogeneity in *Toxoplasma* (Fig. 3). In line with a previous report (22), we can infer multiple instances of transcriptional start site heterogeneity in tachyzoites when we profiled eIF4E1 mRNA binding sites. Transcript isoforms that arise when multiple TSS options are available can differ substantially in their 5’-leader lengths, changing the composition of cis-acting elements that can regulate the translation of the transcript. Determining whether *Toxoplasma* leverages TSS heterogeneity in a developmentally controlled manner should be revisited by assessing eIF4E1-mRNA interactions in tachyzoites and bradyzoites. This strategy has been shown to impart translational selectivity and manage stress adaptation in other systems (30-32).

We also performed the first characterization of eIF4E2. Although homologous within its eIF4E domain and associated with polysomes and stress granules, eIF4E2 surprisingly does not bind the 5’-cap of mRNAs, nor does it associate with the expected eIF4F machinery. In contrast to eIF4E1, depletion of eIF4E2 has no appreciable fitness defect in tachyzoites (Fig. S2). Why eIF4E2 is more abundant than eIF4E1 adds to the mystery of its role in tachyzoites (Fig. 1). Its N- and/or C-terminal extensions may be indicative play a regulatory role in responding to a stress or developmental signal. Ongoing studies are addressing detailed functions of each eIF4E subunit in developing bradyzoites.

### Translation regulation by *Toxoplasma* eIF4F complexes

Mechanisms of eIF4F-based translational regulation vary but can be broadly categorized into two main processes: (i) assembly of paralogous eIF4F complexes and (ii) modulation of eIF4F assembly, both of which can selectively drive translation of subsets of mRNAs. An example of paralogous eIF4F complexes is the selective mRNA interactions of eIF4F and eIFiso4F found in plants, which preferentially interact with different mRNA pools to facilitate translational control (15, 33). Like the usage of paralogous eIF4F complexes in plants or in kinetoplastids (34), we found that eIF4E1 binds two of the three eIF4G orthologs in tachyzoites, raising the possibility of paralogous eIF4F complexes in *Toxoplasma* (Fig. 2). Whether *Toxoplasma*’s eIF4F paralogous complexes differ in function and composition beyond eIF4G subunits remains to be determined. Any transcript preferences exhibited by the paralogous eIF4F complexes could uncover important regulatory features in *Toxoplasma* development that could be leveraged as novel drug targets.

A well-documented example of the modulation of eIF4F assembly mechanism involves the mTORC1-directed translational control of mRNAs with 5’-terminal oligopyrimidine tracks (5’-TOPs). Under the stress of nutrient scarcity in mammals, inhibition of mTORC1 leads to hypo-phosphorylation of the eIF4E binding protein 4E-BP, which in turn increases the affinity of 4E-BP for eIF4E at the expense of eIF4G leading to a reduction in translation (4). This is particularly pronounced for mRNAs that encode 5’-TOPs situated near their TSSs, such as found in ribosomal proteins. Consequently, expression of these proteins and hence ribosome abundance is correlates with nutrient availability (35). Our findings reveal that a similar outcome occurs in *Toxoplasma*. Our data show that eIF4E1 depletion produces a sharp decline in the synthesis of ribosome-associated machinery (Fig. 5). However, we found no evidence that this occurs due to a 5’-TOP sequence motif since the same pyrimidine-rich motif was found independent of the efficiency of mRNA translation. It is important to note that the eIF4E-sensitive mRNAs in *Toxoplasma* tend to have 5’-leaders that are much shorter than the long 5’-leaders commonly seen in this parasite. These results suggest that ribosome abundance in *Toxoplasma* is regulated by an alternative mechanism driven by eIF4F. Furthermore, given that bradyzoites display reduced translation as evidenced by puromycin incorporation (Fig. 6), we posit that a reduction in ribosome content may be an early and key part of the differentiation process and is required for bradyzoite persistence analogous to what occurs in other systems (36).

The reduced eIF4E1 abundance in bradyzoites raises the question: how does translation occur in this life cycle stage? There are several possible non-exclusive explanations. First, reduction of eIF4E1 levels may itself be sufficient to enable the selective translation required for bradyzoite development and persistence as scarcity of the factor may promote selectivity in eIF4E1-mRNA engagement as previously reported (30). A second possibility is that another eIF4E paralog, such as eIF4E2, may play a more predominant role in promoting cap-dependent translation in bradyzoites. Third, a reduction in cap-dependent translation may increase the prevalence of cap-independent translation mechanisms which may play a role in bradyzoite formation. We note that *Toxoplasma* eIF4G3 lacks an identifiable eIF4E-binding motif and resembles the organization of mammalian eIF4G2/DAP5, which promotes cap-independent translation (20). The finding that BFD2/ROCY1 protein binds the 5’-leader of the bradyzoite-driving transcription factor *BFD1* mRNA and is required for the latter’s translational activation (10-12), may indicate that features located within the long 5’-leaders of critical mRNAs are central to promoting bradyzoite formation.

### Concluding remarks

The eIF4F complex is a major regulator of translation initiation linked to nutrient sensing, stress adaptation, cell differentiation, and organismal development (4, 13, 19). Here, we addressed the role of eIF4F in *Toxoplasma* mRNA translation and its contribution to bradyzoite formation. Of the three eIF4E domain-containing proteins encoded in *Toxoplasma*, only eIF4E1 binds to the mRNA 5’-cap and eIF4G subunits in tachyzoites. We conclude that eIF4E1 plays the predominant role in driving canonical cap-dependent translation in this developmental stage. We also discovered that targeted eIF4E1 depletion induced robust and spontaneous bradyzoite differentiation in the absence of exogenous stress, suggesting a critical role in controlling *Toxoplasma* latency.

## Materials and methods

### Cell lines and culture conditions

Human foreskin fibroblasts (ATCC SCRC-1041) were maintained in DMEM (pH 7.4) supplemented with 10% FBS and cultured at 37°C and 5% CO_2_. *Toxoplasma gondii* strains used in this study include RHΔKu80, obtained from Dr. Vern Carruthers (37), and RH::TIR1, and ME49::TIR1, obtained from Dr. Kevin Brown and Dr. David Sibley (25, 38). Tachyzoites were cultured in the same media and conditions as the mammalian host cells. For stress-induced bradyzoite conversion experiments, parasites were cultured at 37°C and ambient CO_2_ in RPMI supplemented with 5% FBS and buffered with 50 mM HEPES pH 8.3 as we have performed in the past (39). The alkaline media was changed daily to maintain the pH.

### Endogenous tagging of parasite genes

All endogenously tagged genes were generated by electroporating a plasmid encoding Cas9 and a guide RNA targeting a region near the gene of interest (GOI)’s stop codon along with a repair template designed to introduce either an epitope tag (HA or MYC) or an HA epitope tag fused with a minimal Auxin Inducible Degron (mAID) into the protein’s C-terminus along with a downstream selection cassette by double homologous recombination. pCAS9 (40) was modified by PCR mutagenesis and repair templates were generated by PCR to include a ∼40 nt homology to the 3’end of the CDS and a region in the GOI’s 3’UTR. Clonal parasites were obtained by serial dilution after maintaining transfected parasite populations in culture media supplemented with selection compounds, 25 µg/ml mycophenolic acid and 50 µg/ml xanthine and/or 2 µM pyrimethamine. The genetically modified locus was validated by PCR of genomic DNA from parental and clonal transgenic parasites to ensure double homologous recombination by profiling both 5’ and 3’ integration sites (data not shown). Primers used for mutagenesis and genotyping are in Table S4.

### Immunofluorescence assays

Fibroblasts were grown to confluency on glass coverslips. Infected monolayers were fixed in 4% paraformaldehyde prior to permeabilization in blocking buffer (1X PBS, 3% BSA, 1% Triton X-100) for 30 min. eIF4E2 proved resistant to staining when fixed in paraformaldehyde; consequently, 100% cold methanol was used to fix monolayers for assays comparing the stress granule formation of the different eIF4E^HA^/PABP^MYC^ lines. Coverslips were incubated with primary antibodies in blocking buffer for 1 h followed by three 5 min washes in PBS. Coverslips were incubated in blocking buffer containing secondary antibodies along with DAPI and *Dolichos biflorus* lectin as appropriate for 1 h. Coverslips were mounted onto glass slides after 3 x 5 min washes. Antibody source and dilutions are in Table S4.

For puromycin labeling, infected monolayers were treated for 10 min with 100 mg/ml cycloheximide (CHX) and/or 10 µg/ml puromycin immediately prior to fixation in paraformaldehyde. Monolayers were treated with 100 mg/ml CHX and/or 500 µM NaAsO_2_ for 1 h immediately prior to fixation to assess stress granule formation.

For replication assays, parasites were inoculated onto confluent monolayers and allowed to invade for 4 h. The media was aspirated and replaced with media containing 500 M 3-indoleacetic acid (IAA) or 0.5% DMSO vehicle. The cultures were allowed to progress for an additional 16 h prior to fixation and staining with DAPI as described above. Replication assays were conducted in triplicate at least two independent times. The weighted average number of parasites per vacuole was assessed for statistical significance by Student’s t-test assuming unequal variances.

### Plaque assays

Confluent monolayers were infected with 500 (RH strain) or 5000 (ME49 strain) parasites. The media was changed 24 h later to either DMEM containing 500 µM IAA or 0.5% DMSO to allow for a pulse of eIF4E^mAID-HA^ depletion. The next day, the infected monolayers were washed with DMEM, and then left in vehicle or IAA supplemented DMEM for the indicated times. For the reactivation assay, the media was changed to alkaline RPMI after a 24 h invasion period and the culture was transferred to an incubator with ambient CO_2_. The alkaline media was changed daily for 6 days to maintain pH before allowing for parasite reactivation under tachyzoite culture conditions for the indicated times. Upon completion, all wells were fixed in cold methanol and stained with crystal violet. All plaque assays were performed with four replicates at least two independent times. Differences in plaque number were assessed for statistical significance with a one-way ANOVA followed by Student’s t-test assuming unequal variances.

### Assessment of 4EGI-1 efficacy

The toxicity of 4EGI-1 (Cayman Chemical) on confluent fibroblasts was assessed in a 96-well plate format using serial dilutions of 4EGI-1 and the alamarBlue assay as per manufacturer’s instructions (Thermo Fisher Scientific). The efficacy against *Toxoplasma* was determined using RH strain parasites expressing β-galactosidase in a 96-well format and measuring conversion the metabolism chlorophenol red-beta-D-galactopyranoside (as previously described (41). Fifty percent maximal effective concentrations (EC_50_) were determined with PRISM.

### SDS-PAGE and western blots

RIPA lysis buffer (50 mM Tris pH 7.4, 150 mM NaCl, 0.1% SDS, 0.5% sodium deoxycholate, 1% NP-40) was supplemented with EDTA-free Halt protease and phosphatase inhibitor cocktail (Sigma) and applied directly to PBS-rinsed *Toxoplasma*-infected monolayers. The lysate was sonicated, clarified briefly by centrifugation, and boiled for 10 min in NuPAGE loading buffer. Since the eIF4G paralogs were prone to degradation under the previously described lysis conditions, a modified lysis procedure was used as follows: infected monolayers with eIF4E1^HA^/eIF4G^MYC^ tachyzoites were scrapped into PBS, pelleted briefly, boiled as a dry pellet for 30 second, then resuspended in 1% SDS prior to sonication.

SDS-PAGE was performed with the NuPAGE system (Invitrogen) on 4-12% Bis-Tris gels in either MOPS or MES running buffer depending on the molecular weight of the proteins to be resolved as per manufacturer instructions. Gels were transferred to nitrocellulose membranes which were then blocked for 30 min in blocking buffer (1X TBST, 5% powdered milk). Blots were incubated overnight in primary antibody, washed 3 x 5 min, incubated for 1 h in secondary antibody, then washed another 3 x 5 min. Antibody source and dilutions are in Table S4.

### Immunoprecipitation and m^7^G resin affinity purification

Parasites were grown as tachyzoites in fibroblasts for two days prior to harvesting. Infected monolayers were scraped, syringe lysed, centrifuged, and the pellet was rinsed in PBS to deplete soluble host cell material. The pellet was lysed in IP buffer (50 mM Tris pH 7.4, 150 mM NaCl, 1 mM MgCl_2_, 0.5% NP-40, 10% glycerol) and clarified by centrifugation. To assess eIF4E-m^7^G interaction, lysate was added to 30 µl m^7^G-functionalized (or control) agarose resin (Jena Bioscience) and incubated at 4°C for 4 h. For immunoprecipitation of epitope-tagged proteins, lysate was added to 25 µl Pierce magnetic α-HA or α-MYC beads (Thermo Fisher Scientific) overnight. In all cases, beads were collected and washed 3 x 5 min in IP buffer. Immunoprecipitations for western blotting were boiled in 2X SDS loading dye. Beads for mass spectrometry analysis were washed an additional three times in PBS then sent to the Indiana University School of Medicine Center for Proteome Analysis for on-bead digestion and mass spectrometry analysis. Control immunoprecipitations were conducted using the untagged parental strain for non-specific binding. All co-immunoprecipitations, including the negative control, were performed in biological duplicates. Proteins that were present in at least two immunoprecipitations, absent in the untagged control, and not localized to secretory organelles as predicted by LOPIT (42) were counted as potential interacting candidates.

### Polysome profiling

Polysome profiling paired with western blot analysis was conducted as previously described (43). Infected monolayers were treated with 100 µg/ml CHX prior to collection in lysis buffer (20 mM Tris pH 7.4, 100 mM NaCl, 5 mM MgCl_2_, 100 µg/ml CHX, 1% Triton X-100). Lysate was clarified by centrifugation then applied atop a 10-50% sucrose gradient and centrifuged at 40,000 rpm in a Beckman SW-41-Ti rotor for 2 h at 4°C. Fractions were collected upon gradient profiling using a Biocomp Fractionator equipped with a Gilson fraction collector. Proteins were enriched from each fraction by TCA precipitation and analyzed by western blot as outlined above.

### eIF4E^HA^ CLIPseq

CLIPseq of eIF4E^HA^ parasites was conducted following the eCLIP protocol (21) with modifications for *Toxoplasma* as previously described (12). Tachyzoites were inoculated onto confluent fibroblast monolayers and grown for two days to increase parasite biomass. Infected monolayers were rinsed with PBS and UV crosslinked (254 nM) twice at 175J/cm^2^. The parasites were released from host cells by syringe lysis and rinsed twice with PBS. Cell pellets were resuspended in 1 ml lysis buffer (50 mM Tris pH7.4, 100 mM NaCl, 1% NP40, 0.1% SDS, 0.5% sodium deoxycholate), treated with 40 units RNase I (Ambion) for 5 min, and immunoprecipitated with Pierce α-HA magnetic beads for 6 h after saving an aliquot for the size-matched input sample. mRNA isolation from immunoprecipitated and input samples after SDS-PAGE and subsequent Illumina library preparation was conducted as previously described (21, 44) for three biological replicates. Sequencing libraries were read at 2 x 150 bp and can be accessed at GSE243203.

Illumina adapters were removed from read 2 with Cutadapt (45) and UMIs were called with UMI-tools (46). Sequencing reads of minimal 25 bp in length were depleted *in silico* by mapping to human and *Toxoplasma* rRNA and tRNA sequences with bowtie2 (47). The remaining reads were mapped to the *Toxoplasma* genome (v.54), obtained from toxodb.org (16), with the STAR aligner (48) using the options: - outFilterMismatchNoverLmax = 0.1; -outFilterScoreMinOverLread = 0.75; - outFilterMatchNminOverLread = 0.75; -alignIntronMin 50; -alignIntronMax 5000. PCR duplicates were removed from mapped reads with UMI-tools (46), and eIF4E^HA^-enriched regions were called using PEAKachu (options -adaptive mode; -min_cluster_expr_frac = 10^-6^, -min_block_overlap = 0.8; -min_max_block_expr = 0.8; -mad_multiplier) = 0; - fc_cutoff = 1; -padj_threshold = 0.05) as implemented through the Galaxy CLIPseq explorer platform (49). eIF4E^HA^-enriched regions were annotated with Homer (50) and their proximity to previously published transcriptional start sites (22) and annotated genes was assessed with the Intersect intervals feature of bedtools (51). Metagene plots were generated with deeptools2 (52).

### eIF4E1^mAID-HA^ RIBOseq

Ribosome profiling of eIF4E1^mAID-HA^ parasites was conducted in three biological replicates per condition as previously described (53). Tachyzoites were grown in confluent fibroblast monolayers to increase parasite biomass before treatment with 0.5% DMSO vehicle or 500 µM IAA for 4 h prior to harvest. The infected monolayers were washed twice with PBS supplemented with 100 µg/ml CHX and host cells, lysed by syringe passage and rinsed an additional two times in PBS + CHX to deplete soluble host cell polysomes. The cell pellet was lysed in 750 µl lysis buffer (20 mM Tris pH 7.4, 100 mM NaCl, 10 mM MgCl_2_, 1% Triton X-100, 25U/ml Turbo DNase I), clarified by brief centrifugation, and an aliquot was saved as an input sample. The lysate was incubated with 100 units RNase I on a nutator for 1 h at 4°C, quenched with 200 units of SUPERaseIN (Ambion), and applied to a sucrose gradient for polysome profiling as outlined above. The monosome fraction was isolated and ribosome protected footprints were isolated by gel extraction. The input sample was subjected to alkaline hydrolysis prior to gel extraction of a size-matched input. The samples were sequentially depleted of human then *Toxoplasma* rRNA sequences by species specific riboPOOLs (siTOOLs Biotech) as per manufacturer’s specifications. Subsequent Illumina library preparation was performed as previously described (53, 54). Sequencing libraries were read at 2 x 150 bp and can be accessed at GSE243206.

Illumina adapters were removed from read 1 with Cutadapt (45) and UMIs were called with UMI-tools (46). Sequencing reads of minimal 20 bp in length were depleted *in silico* by mapping to human and *Toxoplasma* rRNA and tRNA sequences with bowtie2 (47). The remaining reads were mapped to the *Toxoplasma* genome (v.54), obtained from toxodb.org (16), with the STAR aligner (48) using the options: - outFilterMultimapNmax = 1; -outFilterScoreMinOverLread = 0.9; - outFilterMatchNminOverLread = 0.9. PCR duplicates were removed from mapped reads with UMI-tools (46), and differential expression was assessed for protein coding genes with DESeq2 (55) while Riborex (56) was used to determine changes in translational efficiency upon eIF4E1^mAID-HA^ depletion. Gene ontology analysis was conducted using the toxodb.org implementation (16) with a Bonferroni-corrected false discovery rate cutoff of 0.01. Length estimates of *Toxoplasma* 5’-leaders were conducted using annotation data available through toxodb.org where Apollo-annotated models (57) (downloaded August 2022) were used as preferred gene models over the current reference gene models. RAMPAGEseq-determined transcriptional start sites (22) were also used to correct reference gene models in the case where Apollo annotations were unavailable. The 5’ proximal motifs of transcripts were determined with the MEME suite (58) using the RNA setting and selecting for motifs between 4-8 nt in length. FPKM estimates were obtained using Cufflinks (59) setting the maximum intron length to 5 kb.

## Acknowledgements

We would like to thank Dr. Kenny Carlson and Sheree Wek for technical assistance. We thank Dr. Kevin Brown and Dr. David Sibley for TIR1-expressing parasites and Dr. Vern Carruthers for the RHΔKu80ΔHX strain. We thank Dr. Gary Ward and Dr. David Sibley for antibodies to IMC1 and aldolase, respectively. The puromycin antibody was deposited to the DSHB by Jonathan Yewdell. This research was supported by grants from the National Institutes of Health (AI172752 and AI167662 to W.J.S. and R.C.W.) and a postdoctoral fellowship from the American Heart Association (20POST35150019) to M.J.H. R.C.W. is a member of the advisory board in HiberCell, Inc. The funders had no role in study design, data collection and analysis, decision to publish, or preparation of the manuscript.

## Supplemental Figure Legends

**Figure S1. The *Toxoplasma* eIF4E family members retain highly conserved features of the eIF4E domain.** A) Multiple sequence alignment (MUSCLE) of eIF4E domains from multiple model species along with the three *Toxoplasma* orthologs. Motifs responsible for eIF4G-binding and key conserved aromatic residues are highlighted and defined in the color-coded index. B) Phylogram of the multiple sequence alignment. C) Western blot of endogenously tagged eIF4E paralogs. The parental strain, RHΔKu80ΔHX, is denoted as RH in the first lane of each blot. SAG1 was probed as a loading control. Molecular weight markers are indicated in kDa. D) Immunofluorescence microscopy of each HA-tagged eIF4E paralog. DNA is stained with DAPI. Microscopy scale bar = 5 µm.

**Figure S2. eIF4E2 is dispensable for tachyzoite fitness and does not operate within an eIF4F complex.** A) Analysis of eIF4E2^HA^ and PABP^MYC^ sedimentation by polysome profiling of tachyzoites exposed to oxidative stress induced by incubation with 500 µM NaAsO_2_ for 1 h or no treatment. The trace is generated by continuous absorbance at A254 which measures RNA content with major peaks at rRNA subunits, monosome, and heavy polysomes. Fractions of equal volumes were collected from the gradient and proteins were enriched by TCA precipitation and analyzed by western blot. An antibody directed against RPS17 that is suggested to recognize both *Toxoplasma* and human orthologs was included on the blot as a reference marker for protein sedimentation profiles. B) Analysis of eIF4E2 localization to stress granules upon oxidative stress. eIF4E2^HA^ and PABP1^MYC^ were visualized by immunofluorescence microscopy and DNA by DAPI staining. Samples were treated with 500 µM NaAsO_2_ ± 100 µg/ml cycloheximide (CHX) for 1 h prior to fixation. The translation elongation inhibitor CHX was included as a control to prevent stress granule formation. Microscopy scale bar = 2 µm. C) eIF4E2^HA^ interactomes under stressed (y-axis; 500 µM NaAsO_2_ for 1 h) or non-stressed (x-axis) tachyzoites. Affinity purification of eIF4E2^HA^ was conducted by immunoprecipitation with α-HA magnetic beads followed by mass spectrometry identification from two biological replicates. Interacting proteins that were absent in the untagged control immunoprecipitations and not localized to *Toxoplasma* secretory organelles (42) are displayed and colored by functional identity. The average percentage of coverage from each identified protein is displayed. D) A metagene plot demonstrating CLIPseq profiles of eIF4E2^HA^-enriched and size-matched input samples. E) Analysis of eIF4E2^mAID-HA^ depletion over time upon addition of 500 µM IAA by western blot. The blot was probed with α-SAG1 as a loading control. Molecular weights are indicated in kDa. F) Plaque assay of vehicle (0.5% DMSO)-treated parasites, parasites pulsed for 24 h with 500 µM IAA or parasites treated with IAA for 6 days. Mean plaque numbers ± standard deviation were tested for statistical significance by a one-way ANOVA followed by Student’s t-test assuming unequal variances. ns = not significant. G) Analysis of parasite replication after a 16 h treatment with 500 µM IAA. Mean number of parasites per vacuole with standard deviation is shown. Statistical significance of the change between the mean number of parasites per vacuole was determined by Student’s t-test assuming unequal variances. ns = not significant.

**Figure S3. eIF4E1 binds mRNA 5’ caps with low selectivity.** A) eIF4E1 engagement with mRNAs is proportional to mRNA abundance as measured by CLIPseq and RNAseq. B) eIF4E1 engagement with mRNA is proportional to abundance of its translation as measured by CLIPseq and RIBOseq. C) The degree of translation largely correlates to mRNA abundance as measured by RIBOseq and RNAseq. Spearman correlation coefficient is presented on the bottom right of each graph. Discordance can be seen by apicoplast-encoded genes (purple) which are transcribed and translated within the organelle. Genes displaying reduced translational efficiency upon eIF4E1^mAID-HA^ depletion are marked in orange.

**Figure S4. Determination of 4EGI-1 activity against *Toxoplasma* and human fibroblasts.** Estimation of the 50% maximal effective concentration of 4EGI-1 against confluent human foreskin fibroblasts was assessed by alamar blue assay. Estimation of the 50% maximal effective concentration of 4EGI-1 against an RH strain *Toxoplasma* line that expresses a β-galactosidase reporter was determined by measuring the conversion of chlorophenol red-beta-D-galactopyranoside. Error bars represent the standard deviation between the replicates.

**Figure S5. Generation of eIF4E1^mAID-HA^ parasites in the ME49 background.** A) Analysis of eIF4E1^mAID-HA^ depletion by western blot in ME49 parasites upon addition of 500 µM IAA. The blot was probed with α-SAG1 as a loading control. Molecular weights are indicated in kDa. B) Immunofluorescence microscopy of eIF4E^mAID-HA^ ME49 parasites upon treatment with DMSO (vehicle) or IAA for 4 h. DNA is stained with DAPI. Microscopy scale bar = 2 µm.

## Supplemental Table Legends

**Table S1.** eIF4E interacting proteins in tachyzoites as determined by immunoprecipitation followed by mass spectrometry analysis.

**Table S2.** Sites of eIF4E-mRNA interactions in tachyzoites as determined by CLIPseq.

**Table S3.** Differential gene expression at the transcriptional and translational level upon eIF4E1^mAID-HA^ depletion in tachyzoites as determined by RIBOseq with paired RNAseq.

**Table S4.** List of oligonucleotides and antibodies used in this study.

