## Supplementary figures and images for "mRNA cap-binding protein eIF4E1 is a novel regulator of *Toxoplasma gondii* latency"

### Fig. S1

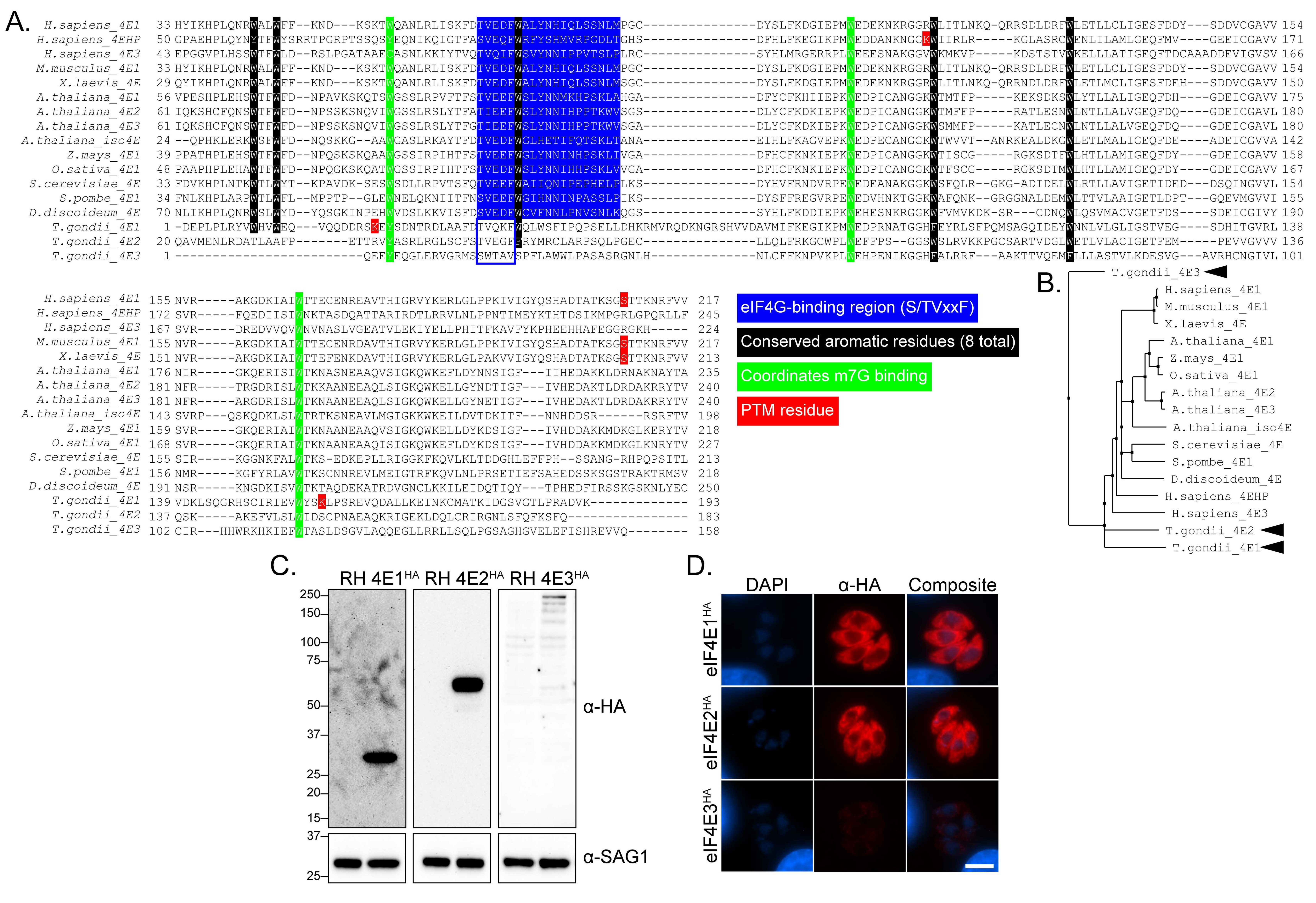

### Fig. S2

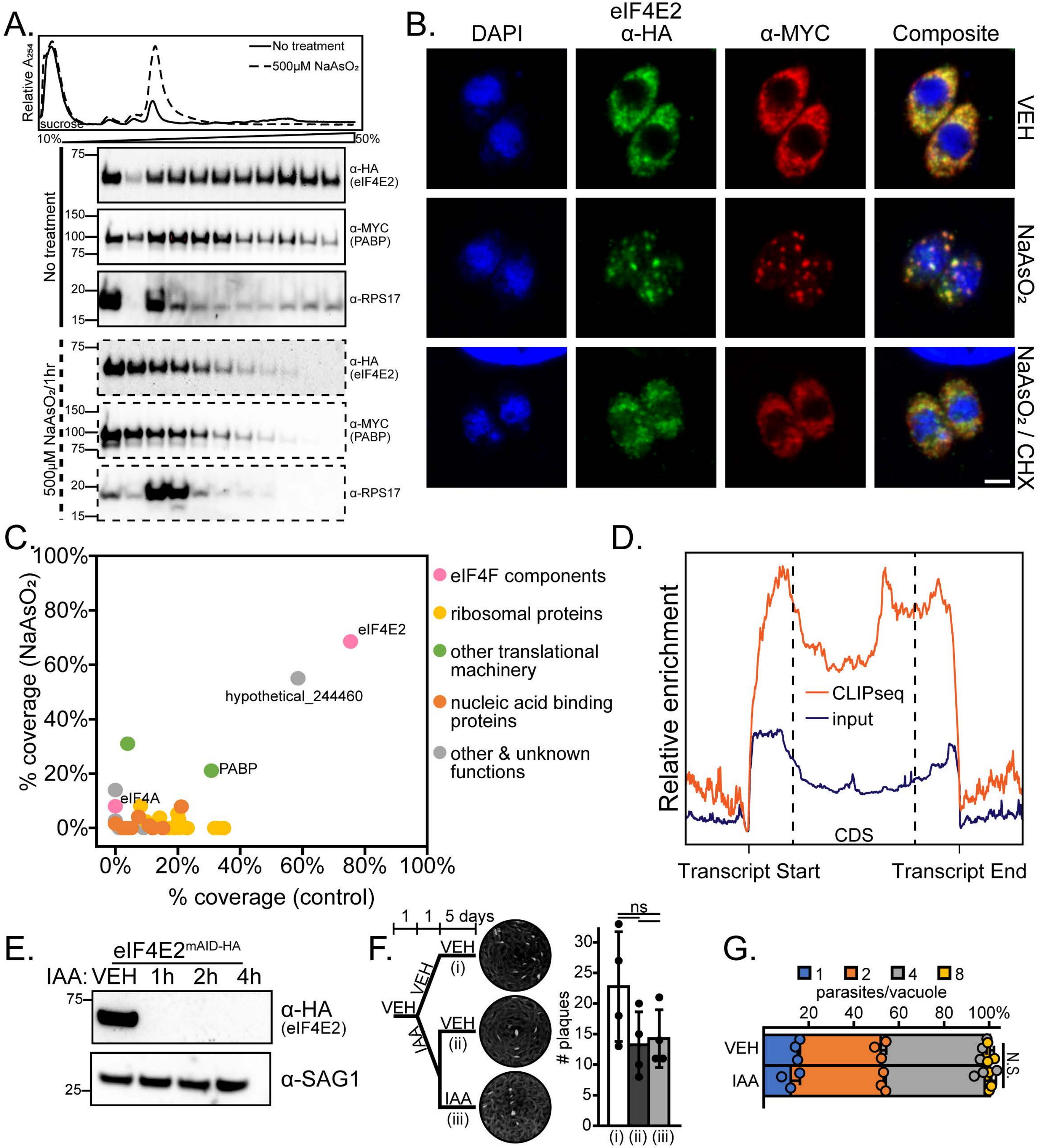

### Fig. S3

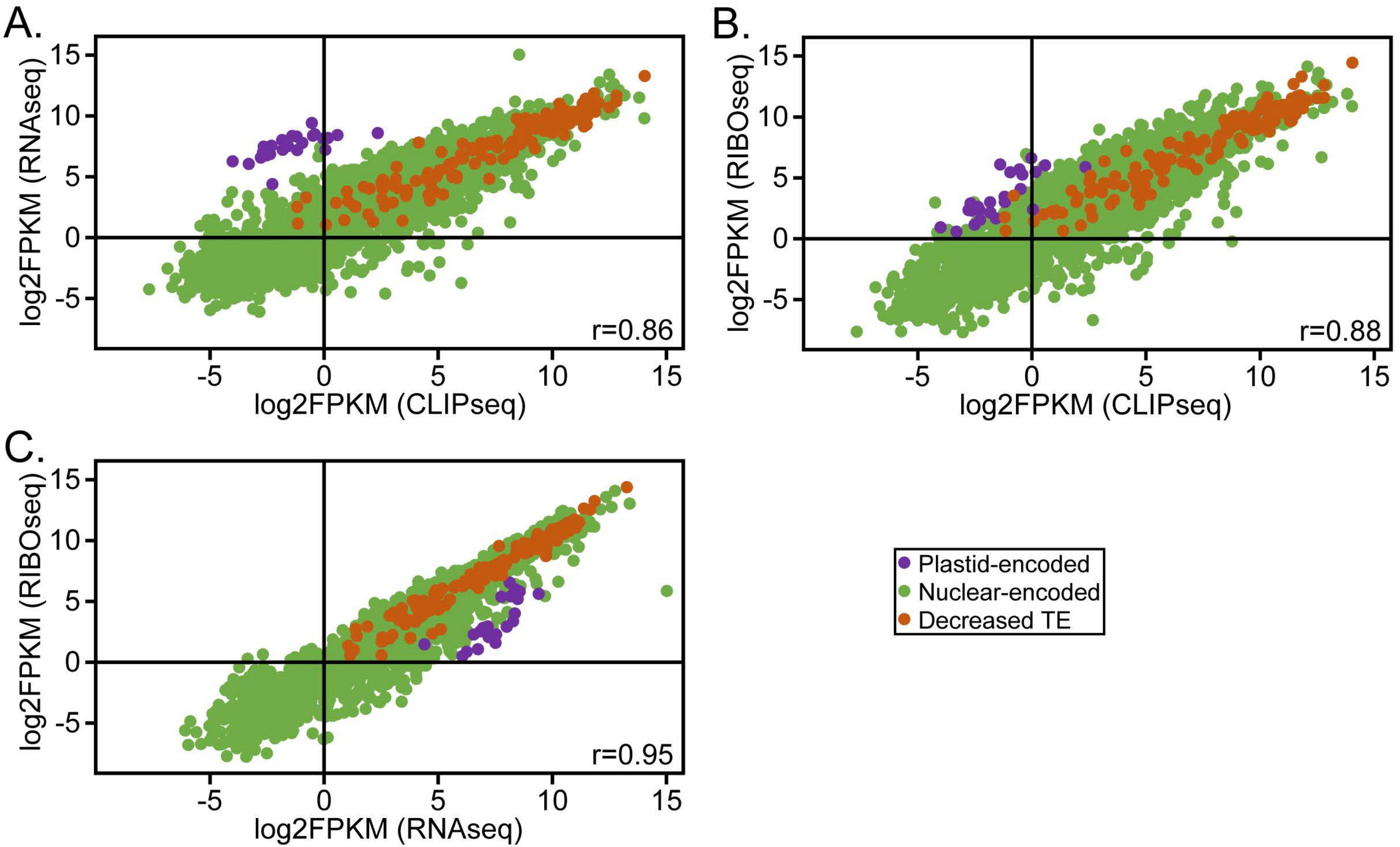

### Fig. S4

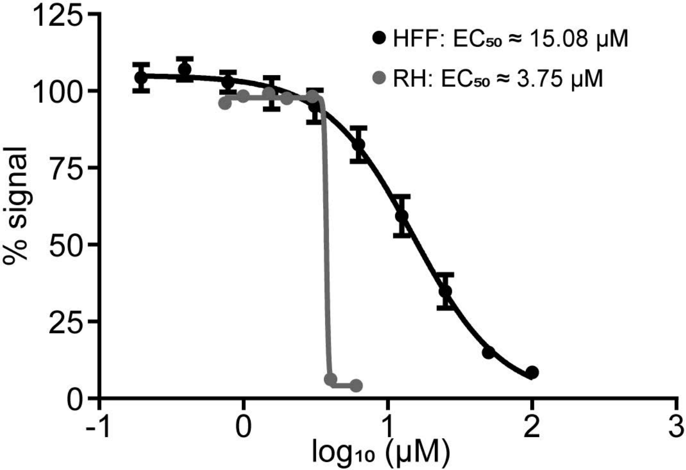

### Fig. S5

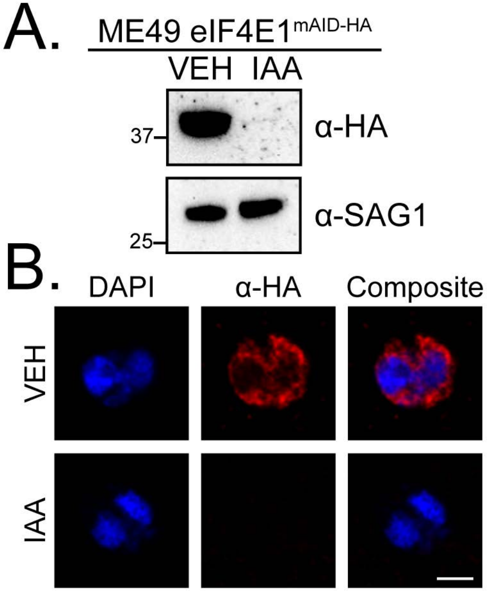
